# Importance of Mobile Genetic Element Immunity in Numerically Abundant *Trichodesmium* Clades

**DOI:** 10.1101/2022.04.20.488952

**Authors:** Eric A. Webb, Noelle A. Held, Yiming Zhao, Elaina Graham, Asa E. Conover, Jake Semones, Michael D. Lee, Yuanyuan Feng, Feixue Fu, Mak A. Saito, David A. Hutchins

## Abstract

The colony-forming cyanobacteria *Trichodesmium* spp. are considered one of the most important nitrogen-fixing genera in the warm, low nutrient, open ocean. Despite this central biogeochemical role, many questions about their evolution, physiology, and trophic interactions remain unanswered. To address these questions, we describe the genetic potential of the genus via significantly improved genomic assemblies of strains *Trichodesmium thiebautii* H94, *Trichodesmium erythraeum* 2175, and 17 new *Trichodesmium* metagenome-assembled genomes (MAGs, >50% complete) from hand-picked, *Trichodesmium* colonies spanning the Atlantic Ocean. Phylogenomics identified ∼four N_2_ fixing clades of *Trichodesmium* across the transect, with *T. thiebautii* dominating the colony-specific reads. Pangenomic analyses showed that all *T. thiebautii* MAGs are enriched in defense mechanisms and encode a vertically inherited Type III-B Clustered Regularly Interspaced Short Palindromic Repeats and associated protein-based immunity system (CRISPR-Cas hereafter). Surprisingly, this CRISPR-Cas system was absent in all *T. erythraeum* genomes and MAGs, vertically inherited by *T. thiebautii*, and correlated with increased signatures of horizontal gene transfer. Multiple lines of evidence indicate that the CRISPR-Cas system is functional in situ: 1. *Trichodesmium* CRISPR spacer sequences with 100% identical hits to field-assembled, putative phage genome fragments were identified, 2. High *Trichodesmium* spacer sequence variability indicating rapid adaptation, and 3. metaproteomic and transcriptomic expression analyses detecting the CRISPR-Cas system components in *Trichodesmium* colonies from the Atlantic and Pacific Oceans. These data suggest that phage or mobile genetic element immunity in *T. thiebautii* could contribute to their success, gene diversity, and numerical dominance over *T. erythraeum* in the oceans, thus warranting further *Trichodesmium* virome investigations.

**Significance statement:** Our work identifies CRISPR-Cas immunity as a phylogenetically distinct, environmentally expressed factor in the speciation of closely related N_2_-fixing *Trichodesmium* clades. These findings suggest that differential phage predation and resistance could be a previously overlooked selective pressure in the genus, potentially leading to the current numerical dominance of *T. thiebautii* over *T. erythraeum* in the oceans. Furthermore, while the currently CO_2_-limited *T. erythraeum* is expected to be a ‘winner’ of anthropogenic climate change, their predicted higher phage sensitivity than *T. thiebautii* could challenge this outcome.

---

Low bioavailable concentrations of nitrogen can limit primary productivity in many oceanic euphotic zones (e.g., (1)). In the warm, oligotrophic open ocean, these low nitrogen concentrations select for nitrogen-fixing organisms that can efficiently convert atmospheric N_2_ to bioavailable NH_4_ or amino acids (2). While our understanding of nitrogen-fixing organisms in the oceans is evolving to include non-autotrophic diazotrophs and other unexpected physiologies (e.g., (3–5)), the filamentous, colony-forming cyanobacterium *Trichodesmium* is still considered a critical oceanic nitrogen fixer (3, 6).

Mariners have known filamentous *Trichodesmium spp*. as ‘sea-saw dust’ for hundreds of years because of the massive surface blooms they can form resembling small, water-suspended wood shavings (6). *Trichodesmium* filaments can aggregate in natural communities, forming 1-4 mm colonies of essentially two morphologies (*i*.*e*., radial tufts or spherical puffs; (7)) that are visible to the naked eye and thus aided these early observations. Fitting with this long-term recognition, botanical scientists defined six morphologically described species as early as the late 1800s (8). Still, oceanographers did not recognize their central role in N_2_ fixation until the 1960s ((2, 3)4/20/2022 2:17:00 PM and references therein). Researchers now know that *Trichodesmium* has a wide distribution in the tropics and subtropics (3, 9) and, even though some appear to have lost N_2_ fixation capabilities (5), the genus is still an essential source of bioavailable N to the oligotrophic oceans (6, 10, 11). Thus, while *Trichodesmium* species names have existed for >100 years, experiments to understand their evolution, genomic potential, and ecological impact are still active research areas.

Members of the *Trichodesmium* genus are closely related. Yet, enrichment strains and field samples can show surprising morphological and physiological character variability (*i*.*e*., pigmentation, cell size, trichome shape, growth rate, N_2_ fixation rate, or colony structure) and abundance differences (e.g., (7, 12–15)). For example, marker gene phylogenetics shows four clades of *Trichodesmium* (7, 12, 16), with the best bootstrap support defining the *Trichodesmium thiebautii* and *Trichodesmium erythraeum*-enriched clades I and III, respectively (7). Additionally, morphological and molecular fieldwork shows that members of these same two clades are commonly observed and that *T. thiebautii-*containing clade I is typically more abundant throughout the water column (e.g., (5, 9, 13, 17, 18)). Thus, while there are six classically defined *Trichodesmium* species of varying pigmentation colors and sizes, *T. thiebautii* typically dominates the in situ community. Despite this recognition, the internal and external factors causing the numerical dominance of *T. thiebautii* are poorly defined.

Herein we used metapangenomics and metaproteomics of enrichment cultures and hand-picked *Trichodesmium* colonies spanning the Atlantic Ocean to define *T. thiebautii* genomic features that help to explain their population dynamics among *Trichodesmium* communities. Our efforts show that predicted mobile genetic element immunity (*i*.*e*., against phage and mobile plasmids) is a defining feature of *T. thiebautii*, as all clade members maintain and express a conserved Type III-B CRISPR-Cas system (19). Many of the *T. thiebautii* CRISPR-Cas spacers matched phage genome fragments assembled from same cruise samples. Thus, differential phage resistance could help explain the current numerical abundance of *T. thiebautii* over *T. erythraeum*. This result also raises questions about the likelihood of *T. erythraeum* displacing *T. thiebautii* as climate changes (14, 20).

## Results and Discussion

### All N_2_-fixing*Trichodesmium* genomes are ‘low’ protein-encoding

Past work with a handful of *Trichodesmium* isolates shows that their genomes are low protein-encoding (*i*.*e*., ∼63%) and enriched in selfish DNA elements (21); however, these observations have never been studied systematically across the genus. To address this, we assembled 17 new *Trichodesmium* MAGs from our hand-picked colonies obtained on the 2018 Atlantic Ocean spanning TriCoLim cruise (**Materials and Methods, Fig. 1**) and compared them to previously published isolate genomes (21) and four MAGs from a Tara Oceans analysis (5). Two of our previously published USC *Trichodesmium* Culture Collection (USCTCC) genomes, *T. thiebautii* H94 and *T. erythraeum* 2175, were significantly improved by MiSeq resequencing the former and Anvi’o refining both using reads from TriCoLim. **Supp. Table 1** lists the final refined CheckM (22) statistics for all MAGs and genomes, data that more than doubles the genomic information available for this important genus.

**Figure 1.**
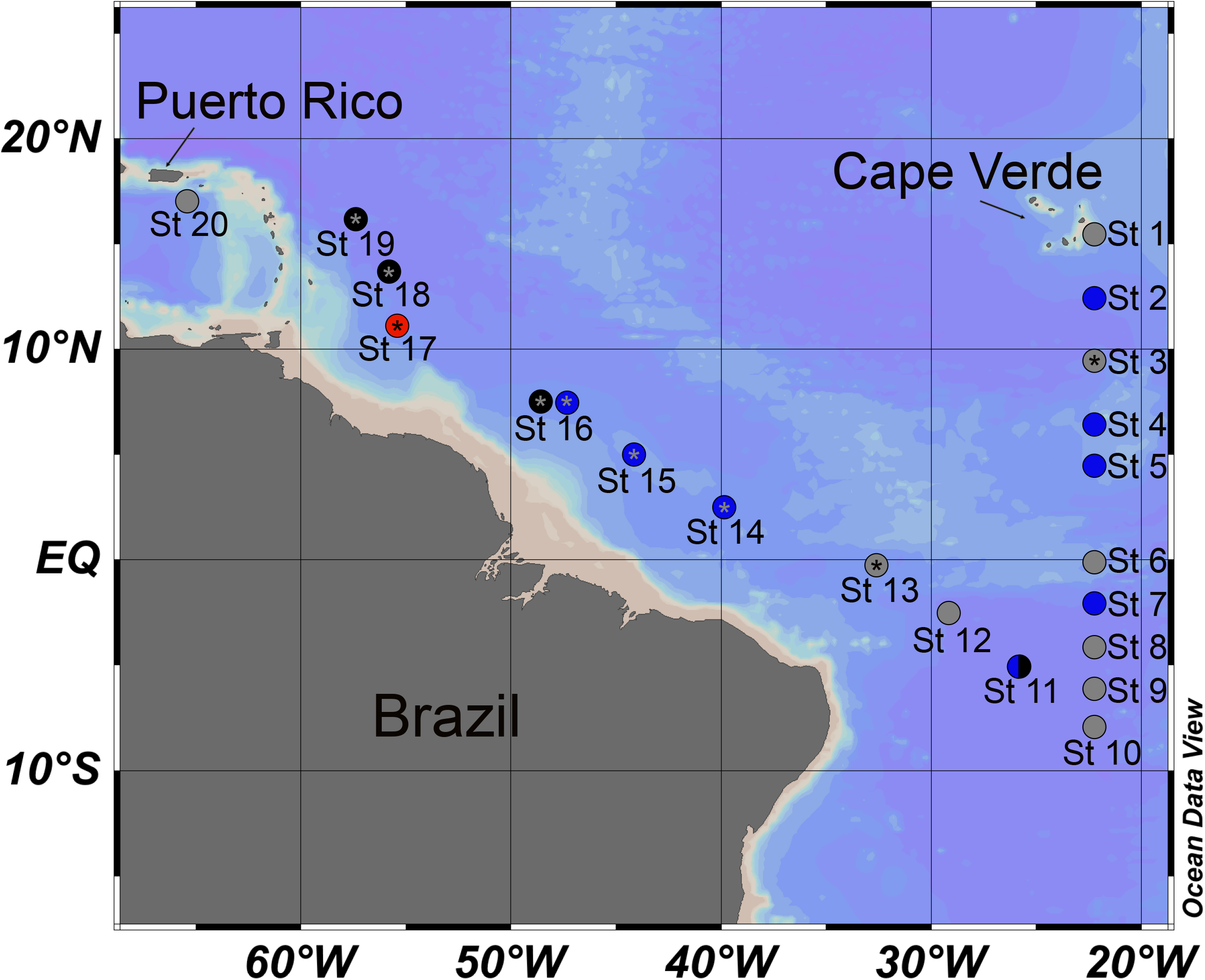
Map of the 2018 Trans-Atlantic TriCoLim Cruise. Color of the station location indicates hand-picked *Trichodesmium* colony morphology, specifically puff (blue), tuft (black), combined (blue and black), not hand-picked (red), no metagenomic data (grey) and metaproteomic data (asterisk).

Similar to the three isolate comparison by Walworth et al. (21), **Supp. Table 2** shows that *Trichodesmium* MAGs >50% complete have low GC% (35 ± 1%) and much lower coding (64 ± 4%) than the bacterial average of ∼90% (23). Because MAG fragmentation and lower completeness scores can significantly change many genome stats, we also calculated these values in higher quality MAGs (*i*.*e*., >80% complete). Our subset obtained a similar coding percentage from above (63 ± 4%) but more consistent values for other parameters. Specifically, *Trichodesmium* genomes are large, with an average length of ∼6.5 ± 0.9 MB, and relatively gene sparse, only encoding for an average of ∼5396 ± 784 proteins.

### There are four N2-fixingclades of *Trichodesmium*

Even though *Trichodesmium* spp. were initially described in the late 1800s, their phylogenetic relationships are still being developed. For example, research using one to three genes has shown that *Trichodesmium* genus members are distributed in two-to-four clades. However, much of the inferred tree topology in those studies was lower support (7, 12, 24). To improve *Trichodesmium* cladistics, we performed phylogenomics using 251 conserved core genes from *Trichodesmium* genomes and MAGs >50% complete. The resulting tree in **Fig. 2A** shows that the N_2_-fixing members of the genus are divided into four major clades. While the tree will likely be improved by including more taxa to resolve subclade structure, it also needs more isolate genomes or MAGs from other oceanic basins (*i*.*e*., only three MAGs were from outside of the Atlantic). That said, while all *Trichodesmium* MAGs are very closely related by average nucleotide identity (>88% ANI), the tree suggests that the *T. thiebautii* assembled from the Atlantic (Clade A) are phylogenomically different from those from other basins (Clade B). However, our read recruiting and metaproteomics (see below) indicate that genomes with high identity to *T. thiebautii* B are also present and active in the Atlantic Ocean, but they did not assemble with high quality from TriCoLim samples (e.g., **Supp Table 1**; St11_bin2_1_1 and St14_bin2_1 are >98% ANI with *T. thiebautii* B H94 and MAG_*Trichodesmium_thiebautii*_Indian). Lastly, while Delmont has shown that some clades of *Trichodesmium* are non-N_2_-fixers (5), BLAST searches of the TriCoLim MAGs with *T. erythraeum* IMS101 *nif* genes confirmed that all of these MAGs likely encode for diazotrophy (*i*.*e*., when missing in a MAG annotation, *nif* genes were consistently found as gene fragments at the end of contigs).

**Figure 2.**
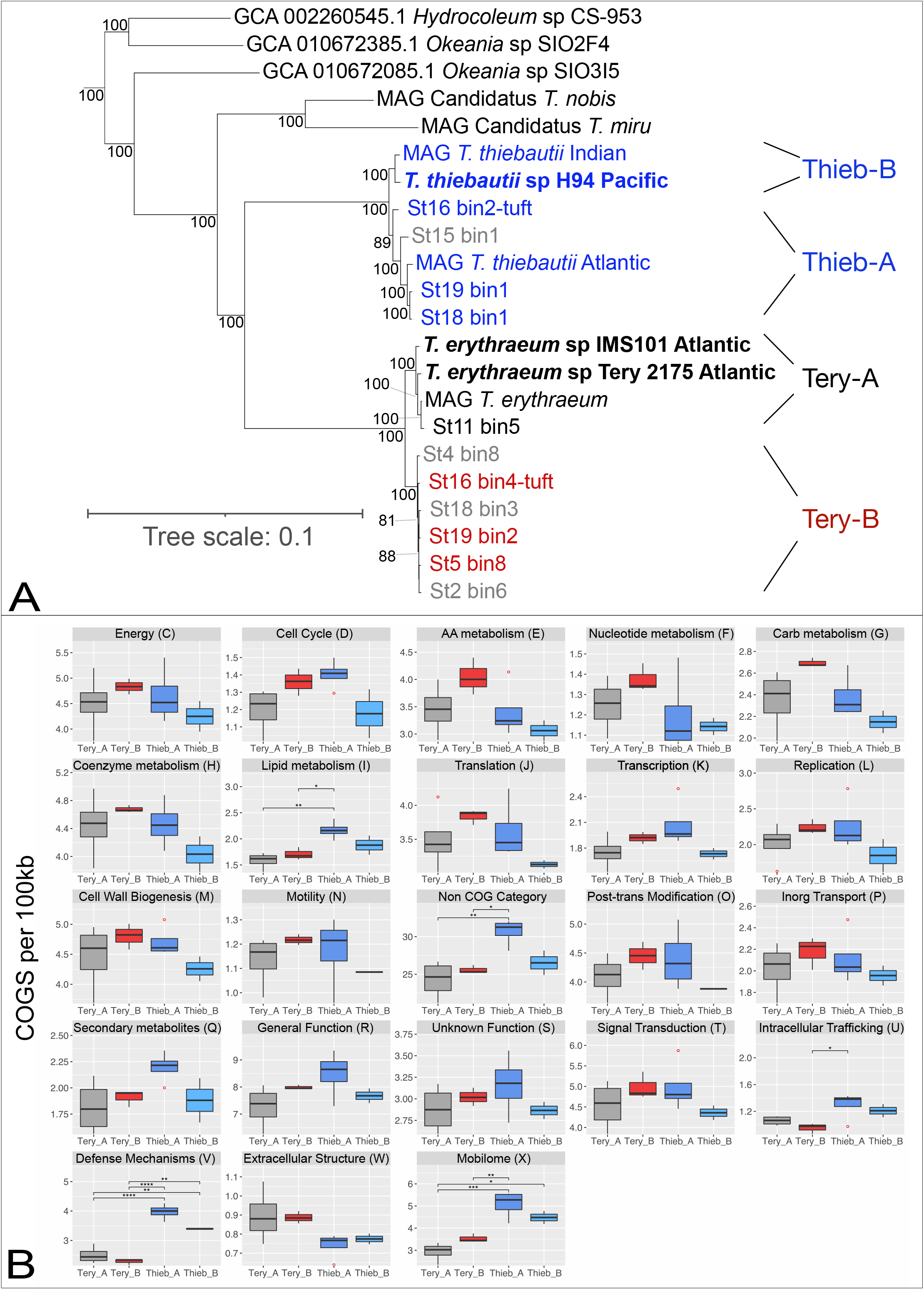
Phylogenomic tree of *Trichodesmium* and nearest relatives and COG enrichment per 100Kb of *Trichodesmium* clades. In panel **A**, TriColim bins are named by station, USCTCC strains are in bold, and names beginning in ‘MAG’ are from (5). *Trichodesmium* MAGs >80% complete are color coded by clade (*i*.*e*., *T. thiebautii* (Blue), *T. erythraeum* A (Black), *T. erythraeum* B (Red), while MAGs <80% are shown in grey. Other *Trichodesmium* MAGs in Supp. Table 1 were excluded from the tree due to low completion values. Panel **B** shows the normalized and summed quantity of COG categories for each MAG per clade. Significantly different categories determined by ANOVA of the mean are denoted above the bracket (p <0.05 =*, <0.01 = **, <0.001 = ***, <0.0001 = ****).

Since there are no published isolate genomes for *T. thiebautii* clade A and *T. erythraeum* clade B, reconciling the past species designations and predicting their physiological and morphological characters was not directly possible. However, we attempted to place these MAGs in context with previously isolated strains by comparing their 16S-23S internal transcribed spacer (ITS) gene sequences with prior *Trichodesmium* enrichment diversity studies (7, 24) via blast. We chose the ITS sequence because past researchers have commonly used this DNA region to probe the diversity and abundance of closely related cyanobacteria (e.g., (25–27)). The isolate ITS hits ranged from 86-100% ID to our MAGs and tracked well with our cladistics. However, the high identity hits (>99.5% ID) allowed us to make three conclusions. **1**. The ITS does not contain enough information to determine if previous *Trichodesmium hildebrandtii, Trichodesmium tenue*, and *Trichodesmium spiralis* isolates are in either *T. thiebautii* clade A or B. **2**. *Trichodesmium contortum* with substantial diameter cells (∼20-30 µm) and bright red coloration is likely a member of *T. erythraeum* clade A. **3**. *T. erythraeum* strains (6-1, 6-2, 6-5) that formed a weak subgroup in a prior analysis (7) are members of *T. erythraeum* clade B. Thus, based on Hynes et al., 2012 we predict that *T. erythraeum* clade B members are phycoerythrobilin-rich red cells ∼6.5-9.5 µm in diameter that can form colonies or loose aggregates. However, more isolate genomes are needed to connect the varied morphologies and pigments observed in *T. thiebautii, T. hildebrandtii*, and *T. tenue* isolates (7) with our phylogenomics. Thus, hereafter we forgo using other *Trichodesmium* species names and simply use the broad clade designations (*i*.*e*., *T. erythraeum* A & B and *T. thiebautii* A & B).

### *Trichodesmium* genomes have many paralogous genes dominated by predicted mobile genetic elements

To understand broad-level genome evolution in the genus, we explored copy number enriched gene families in *Trichodesmium* genomes and MAGs. To accomplish this, we imported each assembly into the Anvio’o pangenomic interface (28, 29), segregated prodigal predicted genes into gene clusters (GCs) with BLAST (30, 31), and annotated these GCs with the 2020 COGs (32) and PFAM (33) datasets. Our results show many paralogous GCs shared by all *Trichodesmium* genomes, with some found in very high copy numbers per clade. Interestingly, each clade’s top ten duplicated GCs are similar but not 100% identical in sequence or copy number (**Supp. Table 3)**. The predicted annotation of these GCs shows that they are enriched (∼78%) in selfish DNA elements like transposases or retrons. For example, GC_00000001 (predicted to be a transposase) is present in 10 of the 13 genomes at 327 copies but is heavily enriched in *T. erythraeum* A MAGs (*i*.*e*., MAG_*Trichodesmium*_*erythraeum* and St11_bin5). One other GC shared by all *Trichodesmium* clades found in high copy numbers was GC_00000002. This GC is annotated as ‘Retron-type reverse transcriptase’ and comprises 342 genes found in as high as 38 copies per genome. Our annotation predicts that this retron is part of a group II self-splicing intron. While the fundamental roles of retrons and group II introns in microbiology are still developing, there is some evidence that they can be involved in bacterial genome re-arrangement or niche adaptation (34–37). Both GC_00000003 (annotated as ‘Transposase|CRISPR-associated protein Csa3, CARF domain (Csa3)’) and GC_00000004 (annotated as ‘Transposase’) are >3-times copy number enriched in *T. thiebautii* clades compared to *T. erythraeum*. Finally, the most abundant duplicated GC in *T. erythraeum* B, GC_00000008 (predicted to be a transposase), is ∼>20 times more abundant in this clade compared to others that maintain it, and it is absent from *T. thiebautii* B. While the factors causing these enrichments/depletions are not defined, these data corroborate a prior finding that *Trichodesmium* genomes are repeat-rich (21) and show that these duplications are commonplace in situ. Furthermore, as seen elsewhere (e.g., (38, 39)), these results suggest that transposition or other related duplication generating processes are important evolutionary mechanisms in *Trichodesmium*.

### *T. thiebautii* MAGs are enriched in specific clusters of orthologous genes (COG) compared to *T. erythraeum*

To begin to understand the selective pressures driving speciation in the genus, we next characterized the genomic potential of *Trichodesmium* in a phylogenomic context. At first glance, the average gene number per genome is greater in *T. thiebautii* than *T. erythraeum* (**Supp. Tables 4-7)**. We used the Anvi’o pangenomic interface (28, 29) described above to characterize these differences, combined with R graphing and statistics. As shown in **Fig. 2B**, many of the COG categories per 100kb in each clade are statistically indistinguishable by ANOVA. However, there are five enriched categories in *T. thiebautii* A, *T. thiebautii* B, or both. These include Lipid Metabolism (I), Intracellular trafficking (U), Defense Mechanisms (V), Mobilome (X), and genes not categorized by COG.

Closer inspection of the COG categories enriched can give more insight into clade-specific niche adaptation. Specifically, the Lipid Metabolism COGs show that *T. thiebautii* A has increased acyl-carrier proteins, many of which appear to be involved in polyketide synthases or annotated with multiple functions. These findings suggest increased secondary metabolite production in this clade. Independent analysis with the secondary metabolite prediction software antiSMASH (40) on the same 13 *Trichodesmium* assemblies also predicted that *T. thiebautii* makes specialized compounds. These include compounds similar to the cancer cell toxin Curacin A (41) and the hydrocarbon 1-heptadecene (42). However, despite the interest in understanding how *Trichodesmium* acquires Fe (e.g., (15, 43, 44)), no putative siderophore producing clusters were defined by antiSMASH. The Intracellular Trafficking COGs enriched in *T. thiebautii* are mostly large repeat-rich proteins (*i*.*e*., annotated as predicted “CHAT domains,” “haemagglutination activity domain,” or “tetratricopeptide repeats”), those implicated in attachment, and related to secretory pathways. However, select BLAST analyses showed that many of these proteins’ closest hits are conserved hypotheticals (typically in other cyanobacterial genomes). Putative transposases or related genes were heavily enriched in the Mobilome, Defense, and non-categorized COG categories. NCBI nr BLAST searches showed that hypothetical proteins were also prevalent in non-categorized COG categories gene clusters. The *T. thiebautii* Defense category was enriched in putative toxin-antitoxin proteins (45–47) antiphage systems defined in (48), and CRISPR-Cas genes (19, 49).

### One-quarter of all *Trichodesmium* MAGs have shared gene clusters

To more closely explore differences between *Trichodesmium* clades, we examined specific gene cluster presence/absence, annotated function (*i*.*e*., via COG function, COG pathway (32), PFAM (33), KOFAM (50) and KEGG (51)) and detection in our TriCoLim reads. This effort allowed us to determine if the functionalities enriched or depleted in each clade in **Fig. 2B** were caused by distinct, new gene clusters, paralogous duplications, or deletions. Additionally, the GC matrix generated by Anvi’o allowed us to characterize the existing *Trichodesmium* pangenome and determine its genome fluidity (52).

Our metapangenomic analysis used one closed *Trichodesmium* genome (*i*.*e*., *T. erythraeum* IMS101; (21)) and twelve MAGs >80% complete and is shown in **Fig. 3**. As seen before (53, 54), our data support the supposition that tuft and puff colony morphology does not correlate with specific clades of *Trichodesmium*, as *T. thiebautii* assemblies dominated the read recruiting regardless of hand-picked sample colony morphology (**Fig 3B**; black heat map). Intra-clade average nucleotide identity (ANI) of the MAGs was very high. Thus, in situ quantification of each was not possible because of likely random read recruiting among high ANI genomes (55). However, since this issue would likely only underestimate the abundances of each clade, we report that *T. thiebautii* MAGs were recruiting at least 1-2 orders of magnitude more reads than *T. erythraeum* from TriCoLim colonies.

**Figure 3.**
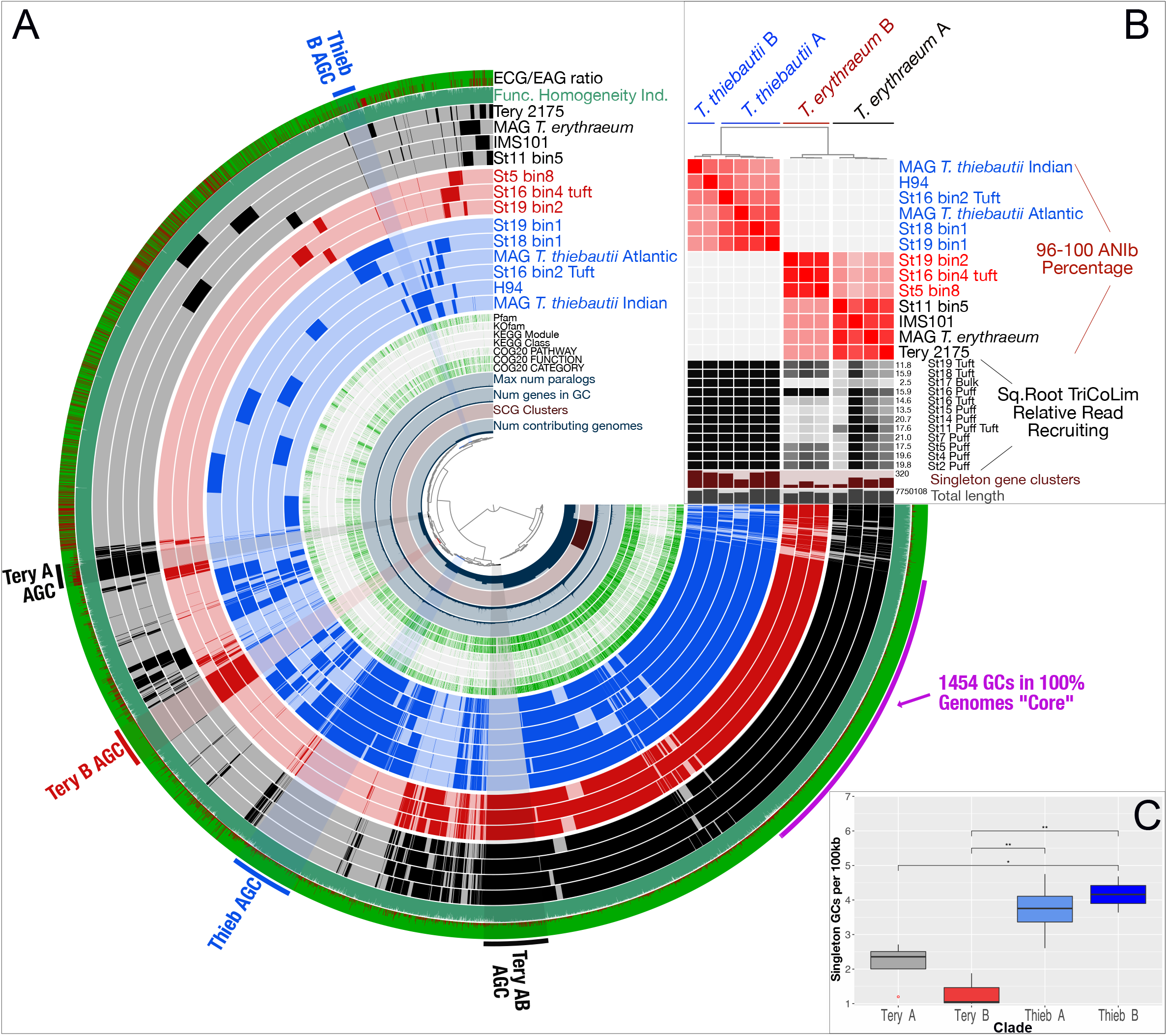
N_2_-fixing *Trichodesmium* Metapangenomic visualization. Panel **A** shows blast-defined conserved gene clusters (GCs) in a MAG as filled colored rings (blue for all *T. thiebautii*, red for *T. erythraeum* B, and black for *T. erythraeum* A). Lighter fill colors indicate that those GCs are missing from that MAG. Singleton GCs (*i*.*e*., appearing in only one MAG) are mostly shown between 9 and 11 o’clock in the phylogram. The innermost rings of the phylogram indicate number of contributing genomes to a GC, single copy genes (SCG), number of genes in a GC, and max number of paralogs. Continuing outwards, if the GC has annotation it is marked in green, while if it does not it is white. The outermost two rings show whether a GC is environmentally core (green) or auxiliary (red; *i*.*e*., the more red the color, the less commonly the GC was observed in TriCoLim reads) and GC homogeneity (*i*.*e*., high homogeneity = all green fill). Clear groupings of clade specific auxiliary gene clusters (AGC) are labeled on the edge of the phylogram. Panel **B** shows ANI clustering at the top and the ANI heat map in red. The black heat map shows square root normalized read recruiting to each MAG from TriCoLim (Blacker bars = higher read recruiting). Panel **C** shows statistical analysis of singleton GCs per clade with ANOVA statistical support shown above the brackets (p <0.05 =*,, 0.01 = **).

Next, we sought to understand the minimum and maximum genetic potential encoded in *Trichodesmium* genomes. Based on the BLAST clustering, there are 1454 blast-derived, single and paralogous GCs in the conservative *Trichodesmium* core found in all genomes (**Fig. 3A)**. Thus, approximately 1/4 of each genome is conserved core gene content. The total pangenome count was 10,054 genes. Pangenome modeling with the R package micropan (56) obtained Heap’s power law alpha estimates of ∼1 for all *Trichodesmium* MAGs together and slightly >1 for *T. thiebautii* and *T. erythraeum* MAGs individually, indicating that these pangenomes are either ‘completely’ sampled with this dataset (*i*.*e*., closed) or slowly growing logarithmically (57).

Others have argued that genome fluidity (□), a metric of genome dissimilarity, is a better method for estimating the likelihood of identifying new genes as more genomes in a group are sequenced (52, 58). We determined the MAG genome fluidity values for all *Trichodesmium* (□ = 0.303 ± 0.10) and the major clades of *T. thiebautii* (□ = 0.24 ± 0.04), and *T. erythraeum* (□ = 0.18 ± 0.03). Strict interpretation of these data suggests a 30% chance of identifying new genes as more *Trichodesmium* genomes are sequenced – again fitting with a growing/open pangenome. While it is important to note that these □ values will likely improve with increased numbers of genomes in each clade (52), the data are consistent with the *T. thiebautii* pangenome being more ‘open’ than *T. erythraeum* and likely experiencing increased horizontal gene transfer (HGT) incorporation rates than the former.

### *Trichodesmium* auxiliary gene content and genomic average nucleotide identity (ANI) recapitulate the phylogenomic signal

While the predicted *Trichodesmium* core N_2_-fixing genome makes up ∼1/4 of the genes, many auxiliary GCs are also detected. As shown in **Fig. 3A**, some auxiliary GCs were only found in one genome (*i*.*e*., singletons), while others associate with specific clades. The environmentally accessory genes (EAGs; *i*.*e*., not found in situ) to environmentally core genes (ECGs; *i*.*e*., found in situ) ratio shown on the outer ring indicates that many, but not all of these auxiliary GC bins, are commonly detected in Atlantic Ocean *Trichodesmium* colonies. Coloring the rings of **Fig. 3A** by phylogenomic group shows that the auxiliary gene content, average nucleotide identity (ANI; **Fig. 3B**), and phylogenomics of core genes (**Fig 2A)** give the same relationships between *Trichodesmium* clades. Additionally, statistical analysis of the singleton genes shows an uneven distribution in the genus, with *T. thiebautii* genomes maintaining significantly more (**Fig. 3C)**. These empirical data are consistent with the genome fluidity results above and suggest mechanisms that increase novel gene content, like horizontal gene transfer, are more common in *T. thiebautii*.

We next took the GCs in each clade-specific bin, highlighted in **Fig. 3A**, to characterize enriched functionalities. The largest groups of clade-specific genes are found in the primary division between *T. thiebautii* AB and *T. erythraeum* AB, where the former shares 313 GCs and the latter shares 315, respectively. Other clear GC groupings were observed in *T. erythraeum* B (198 GCs), *T. erythraeum* A (118 GCs), and *T. thiebautii* B (114 GCs). We performed percentage normalized COG analyses of these conserved GC bins (**Supp. Fig. 1)**, and this effort showed four things: **1**. Non-COG categorized GCs dominate those found in all bins (ranging from ∼44 to 79%), and 18-75% of these non-categorized GCs are hypotheticals based on BLAST searches against NCBI nr (2021-03-01 version), **2**. The Tery-AB bin has much more COG diversity than the similarly sized Thieb bin (315 and 313 GCs, respectively), **3**. Thieb, Thieb-B, and Tery-B bins are enriched in mobilome sequences (∼10% of the bin’s GCs), while in Tery-A and Tery-AB, the mobilome GCs only account for ∼5% of GCs, and **4**. The Thieb bin has a higher percentage of Defense COGs. While our data show that specific CRISPR-related gene duplications are common in *Trichodesmium* MAGs (Duplicated GCs; **Supp. Table 3)**, the Thieb-specific bin is enriched in CRISPR-Cas immunity genes.

### *T. thiebautii* encodes a complete Type III-B CRISPR-Cas system, while *T. erythraeum* does not

To more rigorously characterize and identify the CRISPR-Cas system in *Trichodesmium*, we scanned all assemblies with CRISPRCasTyper (59). This tool aids in the sometimes difficult task of identifying and typing CRISPR arrays and disparate *cas* loci based on the currently defined 44 subtypes and variants (19). The expanded phylogenomic tree in **Fig. 4A** shows a graphical representation of *cas* gene detection and indicates that all *T. thiebautii* assemblies, and those from nearest phylogenomically-defined relatives (*i*.*e*., specific *Okeania* and *Hydrocoleum* MAGs), are predicted to encode CRISPR-Cas systems (e.g., comprised of >15 genes in *T. thiebautii* H94). At the same time, none of the 13 assembled *T. erythraeum* MAGs or enrichment genomes have them. Importantly, since the Joint Genome Institute closed the *T. erythraeum* IMS101 genome (21), the complete absence of a CRISPR-Cas system in this strain supports that they are also missing from *T. erythraeum* MAGs and, not coincidentally in the gaps of a fragmented MAG.

**Figure 4.**
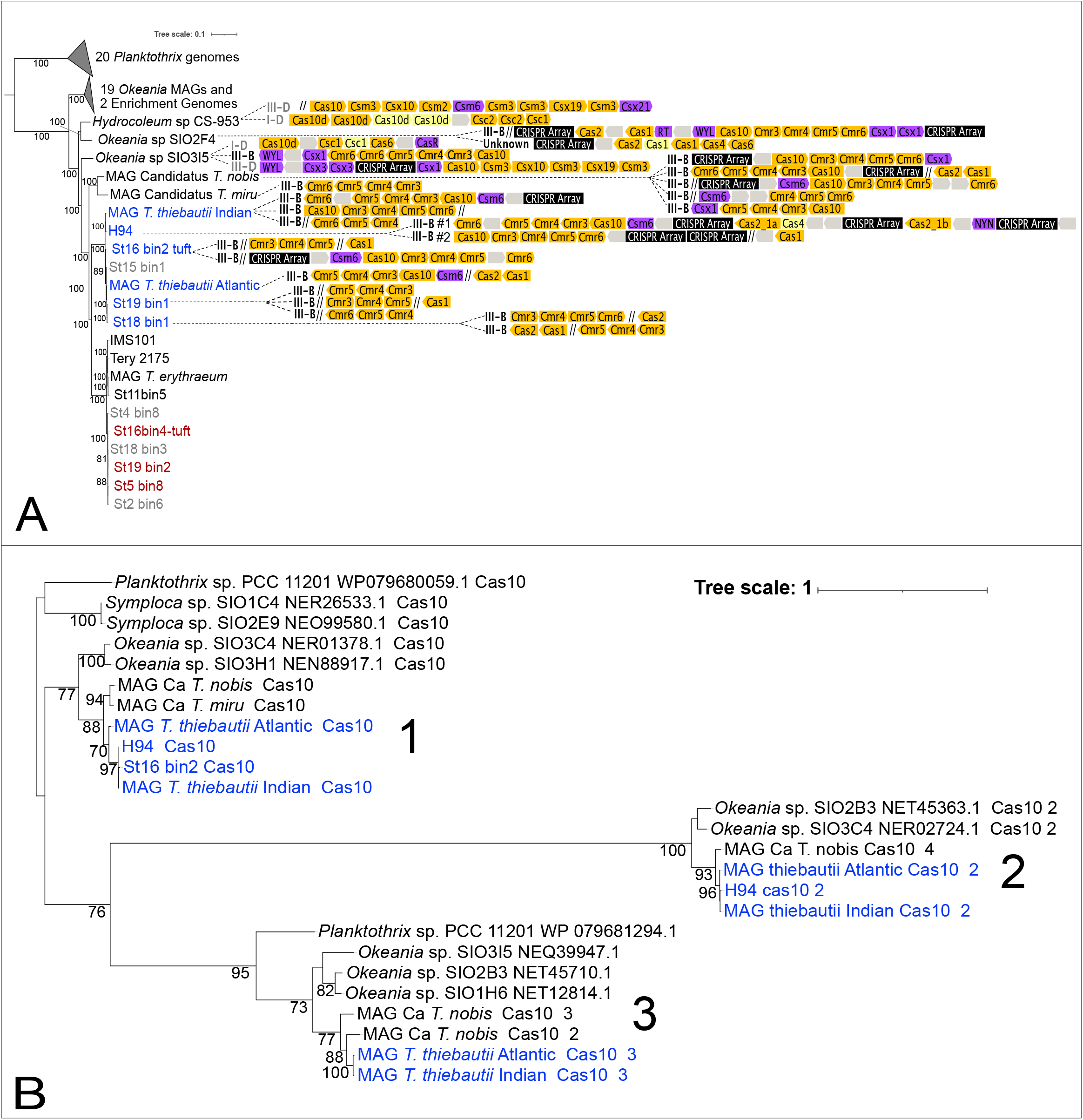
Presence or absence of CRISPR-Cas genes in a *Trichodesmium* MAGs and their nearest relatives phylogenomic context (**A**) and a maximum likelihood tree of their Cas10 protein sequences (**B**). In **A**, the color-coded, directional shapes on the right represent detected Cas genes (yellow), CRISPR arrays (black), Cas accessory genes (purple) and hypothetical genes (grey) that were annotated by as described in the Materials and Methods. Lighter color indicates lower confidence in the annotation. Double-slashes are contig break positions near the annotated CRISPR-Cas systems, indicating that some clusters are fragmented due to breaks in the assemblies. Gene lengths are not drawn to scale. In **B**, the color coding corresponds to *T. thiebautii* (Blue) and other relatives (Black)

As the *cas10* gene is diagnostic for the Type III-B CRISPR predicted to be encoded by *T. thiebautii* and can show significant sequence variation (49), we performed phylogenetic analyses of Cas10 protein sequences to explore the origins of this system in the lineage (**Fig 4A**). The Cas10 maximum likelihood phylogeny shown in **Fig. 4B** suggests two-to-three Type III-B systems in *T. thiebautii*. Additionally, this tree indicates that these systems are likely ancestral because the phylogeny of each of the three distinct sequence clusters is roughly congruent with the phylogenomic signal shown in **Fig. 4A**. However, careful comparison of both trees shows that all three Type III-B Cas10 protein clusters are not conserved in every *T. thiebautii* assembly. ‘Missing’ *cas10* genes lost in assembly gaps or actual deletion of one or more clusters in that MAG could cause this discrepancy. BLASTN searches confirmed that the missing *cas10* genes were present at contig breaks in our *T. thiebautii* MAGs (*i*.*e*., St18_bin1, St19_bin1, and St16_bin2_tuft) corresponding to clusters 1 & 2 in **Fig. 4B**. We could not identify any additional gene fragments for cluster 3 or hits in *T. erythraeum* MAGs. The most straightforward interpretation of these data is that most *T. thiebautii* assemblies do not have cluster 3, and perhaps it is currently disappearing from the *Trichodesmium* pangenome. Fittingly, cluster 3 is undetectable from our best-assembled MAG, *T. thiebautii* H94 isolate genome (566 contigs). Thus, unlike the commonly observed stochastic presence/absence of CRISPR-Cas systems in closely related bacteria (e.g., (60–63), the loss of the III-B CRISPR-Cas system in *T. erythraeum* is phylogenetically constrained and is a defining difference between the major clades of the genus.

Generally speaking, CRISPR-Cas systems protect the cell from mobile genetic elements (MGEs; phage and mobile plasmids) via a sequence-based, targeted genome degradation (60, 64). Many different CRISPR-Cas systems that vary in gene content and recognition molecule (RNA vs. DNA) have been described (19). That said, while all CRISPR-Cas variants appear to provide memory-driven immunity against MGEs, the Type III-B subtype, predicted in numerically abundant *T. thiebautii* clades, requires active RNA transcription for function, can use other CRISPR arrays in addition to its own and provides better protection against phage protospacer mutagenic evasion (65).

Mechanistically, Type III-B CRISPR-Cas systems operate in three steps **1**. Adaptation: recognition and incorporation of transcribed 30-50bp protospacers (*i*.*e*., DNA or RNA sequences of invading MGEs; typically mediated by Cas1 or Cas1-reverse transcriptase (RT) fusion proteins, respectively (66, 67)) into CRISPR arrays as spacer sequence DNA ‘memories’ of past attacks, **2**. Expression: spacer RNAs are expressed as precursor CRISPR RNA (crRNA), and **3**. Interference: sequence-specific crRNAs guides interfere with invading phage or plasmids by the action of the Cas10 protein (49, 60). The absence of a Cas1-RT fusion protein in *T. thiebautii* suggests that the primary adaptation targets for this system are DNA MGEs. In contrast, an HD superfamily nuclease domain in *T. thiebautii* Cas10 proteins indicates that the interference step is likely cleaving both RNA and transcribed DNA (49, 66, 68). Importantly, these DNA spacer sequences also provide ‘fingerprints’ of past MGE attacks that link phage/plasmid sequences with the CRISPR-Cas system containing host (e.g., (69–74)).

### Predicted phage genome fragments assembled from TriCoLim 100% match *T. thiebautii* CRISPR spacers

We next sought to identify predicted MGEs from colony assemblies and determine if they matched *T. thiebautii* spacer sequences. To accomplish this, we asked if putative phage genomes were detectible in our assemblies (*i*.*e*., from enrichment cultures and TriCoLim) using virsorter2 and DRAM-V (75, 76), and we attempted to assemble plasmids using metaplasmidSPAdes (77). While metaplasmidSPAdes identified several putative plasmids in enrichment and field samples (data not shown), none matched any *T. thiebautii* spacers. We also could not detect phage particles/genomes from the enrichment MAGs; however, the TriCoLim assemblies revealed 1000s of putative phage genome fragments with contigs sizes ranging from 1000s to >100kbps (data not shown).

Next, we asked whether these putative phage genomes had sequences matching the *T. thiebautii* spacers using *Trichodesmium* spacer blast searches against the complete DRAM-V predicted phage genomes dataset. This effort identified seven 100% ID hits and 29 more with ANI >90% covering ≥ 93% of the spacer (**Supp. Table 8)**. We conservatively picked the latter ID and coverage level because Type III-B crisper systems can function with mismatches, a feature that requires phage to delete ‘whole’ spacer-protospacer targets from their genomes to escape degradation (78–80). Unfortunately, these spacer-matching putative phage DNA fragments only ranged from 1763 to 5636 bps and were thus too small to identify the phage. All spacer-matching contigs were categorized as virstorter2 category 2 (*i*.*e*., likely phage DNA) and contained many predicted hypothetical viral genes, while one also is expected to encode a transposase (**Supp. Table 9)**. It is noteworthy that many of these putative phage genome fragments were detected multiple times from independently assembled TriCoLim stations (**Supp. Table 9; fastANI groupings)**, suggesting that some consistent phage particles were present across the transect. As past research shows that high phage relatedness selects for CRISPR-Cas systems (60, 81, 82), these data suggest that the *T. thiebautii* CRISPR-Cas system is defending against a relatively conserved phage group.

### *T. thiebautii* CRISPR-Cas systems are expressed in situ

Identifying conserved *Trichodesmium* spacers and putative phage genome fragments suggests that the CRISPR-Cas systems are active in the field. To verify, we screened our TriCoLim *Trichodesmium* colony metaproteomics dataset for in situ Cas protein expression. We used non-identical predicted protein sequences from each *Trichodesmium* genome and MAG in **Fig. 3A** as a protein database to screen environmental *Trichodesmium* colony peptide mass spectral data (83). In this re-analysis, we identified 3498 proteins and 68058 peptides. After binning the detected proteins by clade, *T. thiebautii* proteins were >10x more abundant across the transect than either *T. erythraeum* clades (**Fig. 5a** - Red, Pink, Blue, and Cyan filled colored bars, respectively). This protein detection dataset agrees well with our metagenomic read mapping in **Fig. 3B**, where *T. thiebautii* consistently dominated the sequencing reads. Major metabolic proteins such as the photosystems and nitrogenase were detected in high levels across the transect (**Fig. 5B**) and originated mainly from *T. thiebautii* proteins. Non-COG categorized proteins were 50x more abundant across the transect than even these core metabolic functions, consistent with this being one of the most enriched categories in the *T. thiebautii* assemblies. Additionally, the in situ expression of these non-categorized proteins suggests that they are required for environmental growth and highlights the importance of characterizing them further.

**Figure 5.**
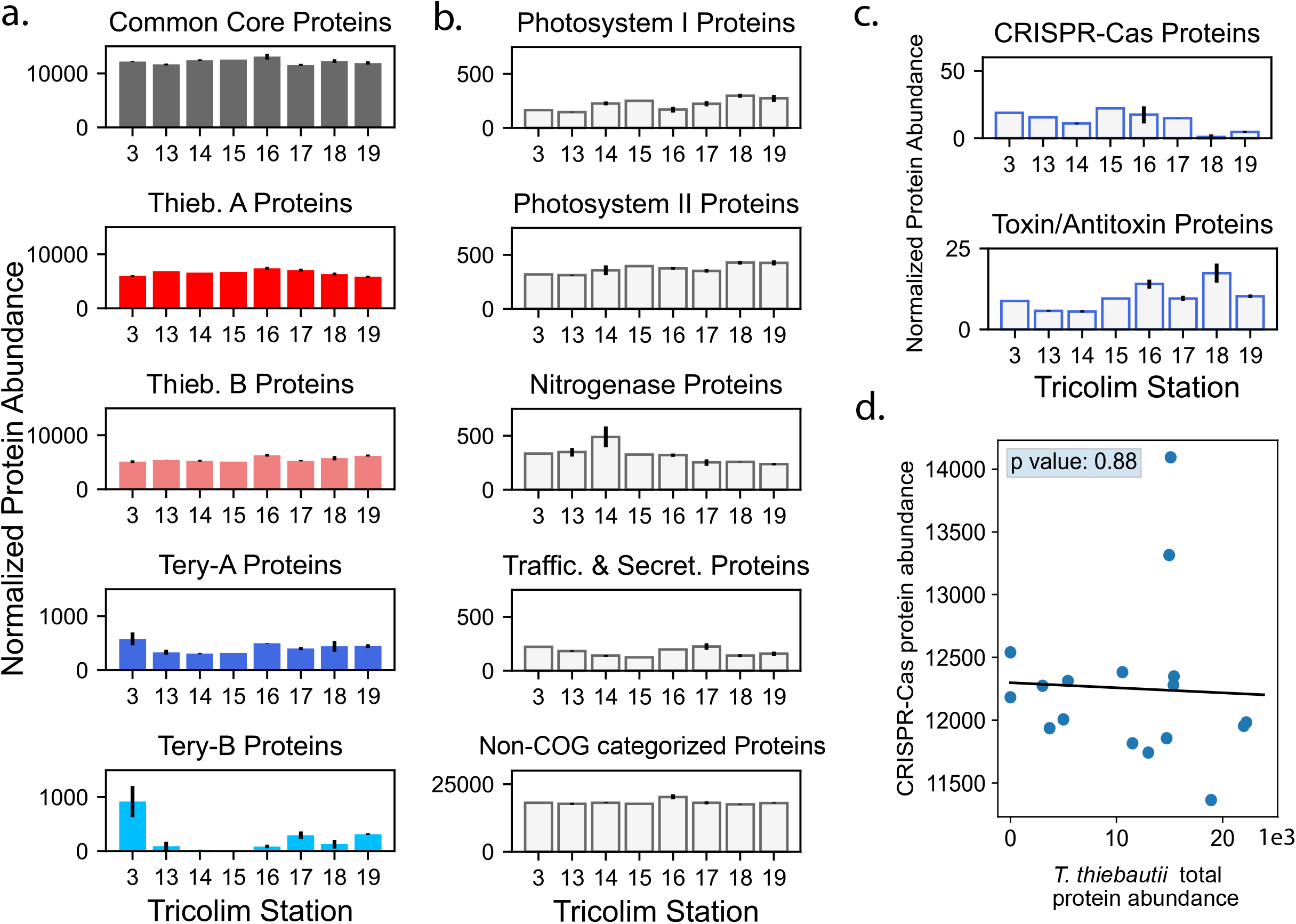
*Trichodesmium* protein abundances across the TriCoLim transect (**A**) Protein abundance data sorted into *Trichodesmium* phylogenetic groups. Proteins were normalized across each sample, then sorted into the respective phylogenetic group and summed. Error bars indicate the standard deviation of the averaged biological replicates. Quantitation is displayed as normalized spectral counts (see Methods). Core and *T. thiebautii* proteins are much more abundant than those derived from *T. erythraeum*. (**B**) Protein abundance data sorted by biological function. Again, proteins were normalized across each sample, then sorted by COG function and summed. Selected, highly abundant functions are shown. (**C**) Summed normalized protein data for CRISPR-Cas and toxin/antitoxin proteins across the TriCoLim transect. (**D**) Nonsignificant correlation of CRISPR-Cas protein abundances versus *T. thiebautii* total protein abundance (p = 0.88).

Proteins involved in cellular defense, including toxin/antitoxin proteins (*i*.*e*., the toxin components of RelE and MazEF and the antitoxin component of ParD) and the CRISPR-Cas system, were identified in appreciable abundance across the transect (**Fig. 5C**). The CRISPR-Cas proteins did not correlate with total *T. thiebautii* protein abundance, suggesting that the former are not constitutively expressed (*i*.*e*., as a function of biomass) and are instead under some regulatory control. The CRISPR proteins identified included Cas10 and Cas7, and their phylogenetic assignment at the peptide level corresponded to *T. thiebautii* species and were assigned Cas10 clusters 1 & 2 from **Fig. 4B**. Specifically, we identified peptides that were identical to those in *T. thiebautii* MAGS H94, St18_bin1, St16_bin2, and MAG *T. thiebautii* Indian Ocean, indicating that these species were contributing to CRISPR-Cas protein production (**Supp. Table 12**). These data also show that Thieb-B clade members are present and active in the Atlantic Ocean.

Research shows that CRISPR-Cas adaptation (*i*.*e*., protospacer incorporation into spacer arrays) requires Cas1 or Cas2 to respond to new MGE threats (19, 67). Thus the absence of these proteins in our metaproteome could suggest that the *T. thiebautii* CRISPR-Cas system is not actively adapting to new phage and is perhaps performing alternative functions in the cell independent of viral immunity (84, 85). Three observations argue against this supposition. **1**. Self-targeting spacers (*i*.*e*., matching alternative sites in the MAG) were not identified, suggesting that interference-based gene regulation is not occurring (e.g., (84–86)). **2**. Most of the spacers detected in each MAG are distinct from those in other MAGs, suggesting that ‘rapid’ adaptation occurs in *T. thiebautii* (69, 87), **3**. Read recruiting from Pacific Ocean *Trichodesmium* community data collected by others (88) shows that all annotated *T. thiebautii* H94 *cas* genes are expressed (including *cas1* and *cas2*), and they appear to have diel periodicity (**Supp. Fig. 2)**. Thus, while we cannot exclude alternative CRISPR functions in *T. thiebautii*, our data strongly suggest CRISPR-Cas mediated phage immunity is commonplace in the clade.

## Conclusions

Herein we show that all N_2_-fixing *Trichodesmium* genomes are large, low protein-coding, and repeat-rich, with *T. thiebautii* having increased singleton gene clusters. Additionally, the conserved maintenance of a functional CRISPR-Cas system in *T. thiebautii* is a defining speciation difference between the major clades of *Trichodesmium* and is likely an important factor in their numerical dominance over *T. erythraeum*. Our findings also raise questions about future oceanic N_2_ fixation – will *T. erythraeum* be a climate change winner as predicted (14, 20) or will increased phage infectivity reduce their future expansion?

The combination of singleton gene enrichment and conserved CRISPR-Cas systems in *T. thiebautii* suggests that immunity allows the recipients of transduced genes to survive and thereby increase their genetic diversity (as noted elsewhere in other systems (89, 90)). Interestingly, CRISPRs are relatively rare in marine systems, and fittingly the numerically dominant planktonic bacteria (e.g., *Prochlorococcus, Synechococcus*, and *Pelagibacter*) do not have CRISPR-Cas systems. Furthermore, many well-described CRISPR-Cas systems are biogeographically confined (*i*.*e*., hot springs) or human health-related. Because of these issues, we are only just beginning to address how MGE selection maintains CRISPR-Cas systems in global populations (81). Thus, *Trichodesmium* colonies represent a globally distributed ‘ecosystem’ to further study MGE-CRISPR-Cas function in a biogeochemically crucial context.

## Materials and Methods

### *Trichodesmium* colony collection

*Trichodesmium* colonies were collected with a hand-towed (∼150 ft of line) 130-µm Sea Gear plankton net on February 8^th^ thru March 11^th^, 2018, during the R/V Atlantis TriCoLim cruise (AT39-05) that transected from the Cape Verde Islands to Puerto Rico (**Fig. 1)**. Colonies were rapidly removed from the cod end and picked separately into tuft and puff morphologies with sterile plastic disposable Pasteur pipettes into sterilized seawater 50 ml Falcon tubes. These morphology-segregated samples were sequentially washed two times, gently filtered down onto 5-µm polycarbonate membranes, and rapidly frozen in liquid N_2_ until processing back at the laboratory.

### DNA Isolation and Sequencing

High-quality DNA was isolated from ∼50 frozen colonies per station via Qiagen DNeasy Powersoil Kit (Germantown, MD) using the manufacturer’s protocol with the following exceptions. Frozen colony samples were rapidly transferred to bead beating tubes with the 5-µm filter unwrapped around the inside of the tube, rendering the biomass containing surface available to the beads. DNA quality and quantity was determined via NanoDrop UV-Vis spectrophotometer and Qubit Fluorometer, respectively (ThermoFisher; Waltham, MA). Samples with lower [DNA] and/or quality were cleaned up with the Qiagen PowerClean Pro DNA clean up kit (Germantown, MD). DNA from twelve TriCoLim samples were 150 PE Illumina sequenced by Novogene (Sacramento, CA) to a final depth of 25 Gbps. DNA was isolated from frozen *T. thiebautii* H94 samples using the same protocol as above and was sequenced via 250PE Illumina MiSeq at the USC Epigenome Center (1.8 Gbps total) because the original assembly in (21) was poor quality. Raw reads are available at NCBI’s SRA under the BioProject PRJNA828267.

### Isolate and Field MAG Assembly

Both the field samples and *T. thiebautii* H94 reads were run through similar assembly pipelines, but the latter was assembled on KBase (https://www.kbase.us), while the former were on a Linux server. The quality of reads was checked with FastQC v0.11.2 (91) and trimmed to enhance stats using Trimmomatic v0.33 (92). MAGs were assembled de novo using metaSPAdes v3.12.0 (93) for H94 and MEGAHIT v1.2.6 (94) for the TriCoLim samples. Binning of contigs was performed via MaxBin2 v2.2.4, quality was checked with CheckM v1.1.3 (22), and phylogenetic placement of the MAGs was determined with GTDB-tk v1.3.0 (95). Field MAGs were dereplicated using fastANI (96) with a cutoff of 98.5% ID. Dereplicated bins >50% CheckM complete were hand refined in Anvi’o v7 (28, 29) until contamination level was below 5%. *T. erythraeum* strain 2175 was downloaded from NCBI and was hand-refined in Anvi’o (28, 29) to remove contaminating contigs using the TriCoLim reads, and its final genome stats were determined with CheckM (22).

### Phylogenomics

Higher quality *Trichodesmium* MAGs (>50% complete; **Supp. Table 1**) and nearest relative genomes downloaded from the NCBI Assembly page were run through the program GToTree v1.6.12 to define a initial guide tree based on 251 cyanobacterial core protein Hidden Markov Models (97, 98). Alignment and partition files from GToTree were piped to IQtree v2.1.4-beta in ModelFinder optimality mode with 1000 ultrafast bootstraps to generate the phylogenomic tree (99). The tree was visualized and edited in the interactive Tree of Life (ITol; (100)).

### Metapangenomics

The metapangenomic pipeline in Anvi’o was used to characterize shared and distinct gene clusters (GCs) in the MAGs and determine if these GCs were represented in the TriCoLim reads (28, 29). Briefly, this pipeline creates a contig database for all MAGs that was then annotated with prodigal (31), COGs (32), PFAMS (33), KOFAM (50) and KEGG (51). Reads were recruited to contigs in the Anvi’o database with TriCoLim read samples using bowtie2 v 2.4.1 (101), matching Sequence Alignment/Map (SAMs) were converted to binary alignment maps (BAMs) with SAMtools v1.11 (102), and BAMs were profiled across the TriCoLim read sets using Anvi’o to determine environmental auxiliary and environmental core genes (EAG and ECG, respectively). COG categories per 100kb was determined by exporting the annotation from Anvi’o, determining the COG categories per MAG, summing those results per clade, and then analyzing and graphing in R v 4.0.3 (2020-10-10) with R Studio v1.4.1103 (103). Differences between COG category counts per clades were tested for statistical significance using ANOVA in the R package rstatix (104). The same general pipeline was used to determine single gene copy per clade and toxin:antitoxin GCs per clade. The Anvio summary data was converting into a matrix via the scripts in (105) and used with the R package micropan (56) to generate Heaps’ law alpha value and genome fluidity estimates.

### CRISPR-Cas Analyses

CRISPRCasTyper (59) was used for the annotation of CRISPR-Cas systems’ subtypes, associated *cas* genes, and CRISPR arrays for all genome assemblies of *Trichodesmium spp*. and MAGs from their nearest relatives (*i*.*e*., *Okeania sp*. SIO2F4, *Okeania sp*. SIO3I5, and *Hydrocoleum sp*. CS-963). The raw output from this program identified CRISPR-Cas systems in multiple contigs in each draft genome and is shown in a phylogenomic context in **Fig. 4A**. Because many of the MAGs were fragmented, CRISPR-Cas system portions on other contigs are shown with double slashes if (1) the pieces were found on the edges of their located contigs, and (2) the associated *cas* genes are still predicted to be part of the subtype III-B, I-D, or III-D systems defined in (19). Additional annotations for accessory genes (purple) and hypothetical genes (grey) were determined by CRISPRCasTyper, CRISPRCasFinder, and BLAST (30, 59, 106, 107).

We used clustal in the program Geneious Prime (Biomatters, San Diego, CA) to align Cas10 protein sequences from all genomes in **Fig. 4A**, and RaxML v8 to generate the maximum likelihood phylogenetic tree with 100 bootstraps (108).

### Virome Assembly

We screened for putative phage genome, prophage or plasmid fragments in TriCoLim and enrichment culture assemblies using the virsorter2, DRAM-V, and checkV pipelines for viruses and metaplasmidSPAdes for plasmids (75–77, 109). The contig sequences supplied from these efforts were used to generate custom blast databases (107) that were subsequently screened with *Trichodesmium* spacers defined above. No hits were defined in the plasmid database, while many high-quality spacer hits were found with DRAM-V putative phage sequences (**Supp. Table 8**). We used FastANI on contigs matching *Trichodesmium* spacers to group them based on >98.1% identity and show that very similar sequences assembled from independent stations (**Supp. Table 9**).

### Proteome analysis of *Trichodesmium* enriched field samples

The raw proteome spectra for this manuscript were collected for a prior work (83) and newly analyzed for this study. Briefly, proteome samples were acquired by gentle hand picking from the phytoplankton nets, rinsed thrice in 0.2-µm filtered trace-metal-clean surface seawater and then decanted onto 0.2-µm supor filters and immediately frozen at -80ºC until protein extraction. Proteins were extracted using a detergent based method and digested while embedded in a polyacrylamide gel, as previously described (83, 110, 111). The global metaproteomes were analyzed by online nanoflow two-dimension active modulation reversed phase-reversed phase liquid chromatography mass spectrometry using a Thermo Orbitrap Fusion instrument (83, 112).

Raw spectra were searched with the Sequest algorithm implemented in Proteome Discoverer 2.2 using a custom-built genomic database consisting of the pan genome of *Trichodesmium*, plus *Trichodesmium* MAGs reported in this study. To avoid inflation of the sequence database and later misinterpretation of phylogenetic signals, only one version of any identical/redundant protein sequences were included in the database, with the possible phylogenetic attributions for the redundant proteins noted in downstream phylogenetic analyses. Sequest mass tolerances were set to +/-10ppm (parent) and +/-0.6 Dalton (fragment). Fixed Cysteine modification of +57.022, and variable N-terminal acetylation of + 42 and methionine modification of +16 were included. Protein identifications were made with Peptide Prophet in Scaffold (Proteome Software) at the 80% peptide threshold minimum, resulting in an estimated peptide FDR of 1.5% and an estimated protein FDR of 0.0%. Relative protein abundances are reported as normalized total exclusive spectral counts, so only spectra corresponding to a specific peptide for a given protein were considered. This avoids the problem of overlapping peptides in the phylogenetic analysis. The values are normalized to total spectral counts identified in each sample. The general workflow for assigning phylogenetic specificity was as follows: if an identified protein was already a member of the *Trichodesmium* core genome, the core designation was retained; for remaining proteins, if all peptide evidence for that protein was present in both *T. erythraeum* and *T. thiebautii* MAGs, the protein was also considered to be “core.” Otherwise, the peptide was assigned to its corresponding MAG phylogeny. Thus, only peptides specific to a given *Trichodesmium* species was considered for that species. The peptides identified for the CRISPR-Cas proteins were further checked using the Metatryp 2.0 tool (www.metatryp.whoi.edu) (113) to ensure phylogenetic specificity of the signals, i.e. to the *T. thiebautii* MAGS specifically mentioned in the text.

The mass spectrometry proteomics data were originally deposited in the ProteomeXchange Consortium via the PRIDE partner repository with the identifier PXD016225 and can be accessed at https://doi.org/10.6019/PXD016225 (114, 115). The data is also available at BCO-DMO (https://www.bco-dmo.org/dataset/787078). The newly searched protein identifications are furthermore provided as **Supp. Tables 10-12**.

### Transcriptome read recruiting

*Trichodesmium* colony metatranscriptomes were downloaded from NCBI SRA (projects PRJNA381915 and PRJNA374879) and mapped against all *cas* genes from H94 and *cas10* from MAG *T. thiebautii* Indian Ocean that is a representative of cluster 3 in **Fig. 4B**. Read quality was checked with FastQC v0.11.2 (91), trimmed with Trimmomatic v0.33 (92), recruited to cas genes with Bowtie v 2.4.1, converted to BAMs with Samtools v1.11, and profiled in Anvi’o v7.0. Average read depth values were normalized with the constitutive *Trichodesmium* gene *rotA*, as in (116). Gene targets and results are summarized in **Supp. Tables 13-15**, and the data were visualized in R v 4.0.3 (2020-10-10) with R Studio v1.4.1103 (103) and shown in **Supp. Fig 2**.

## Acknowledgments

We would like to thank the captain and crew of the RV Atlantis for their essential role in sample collection and providing a safe and efficient platform for marine microbiology. This work was funded by NSF grants OCE 1657757 and OCE 1851222 to DAH, FXF, and EAW, OCE 1850719 to MAS, and by discretionary USC funds to EAW.

