## Supplemental Figures for "Importance of Mobile Genetic Element Immunity in Numerically Abundant *Trichodesmium* Clades"

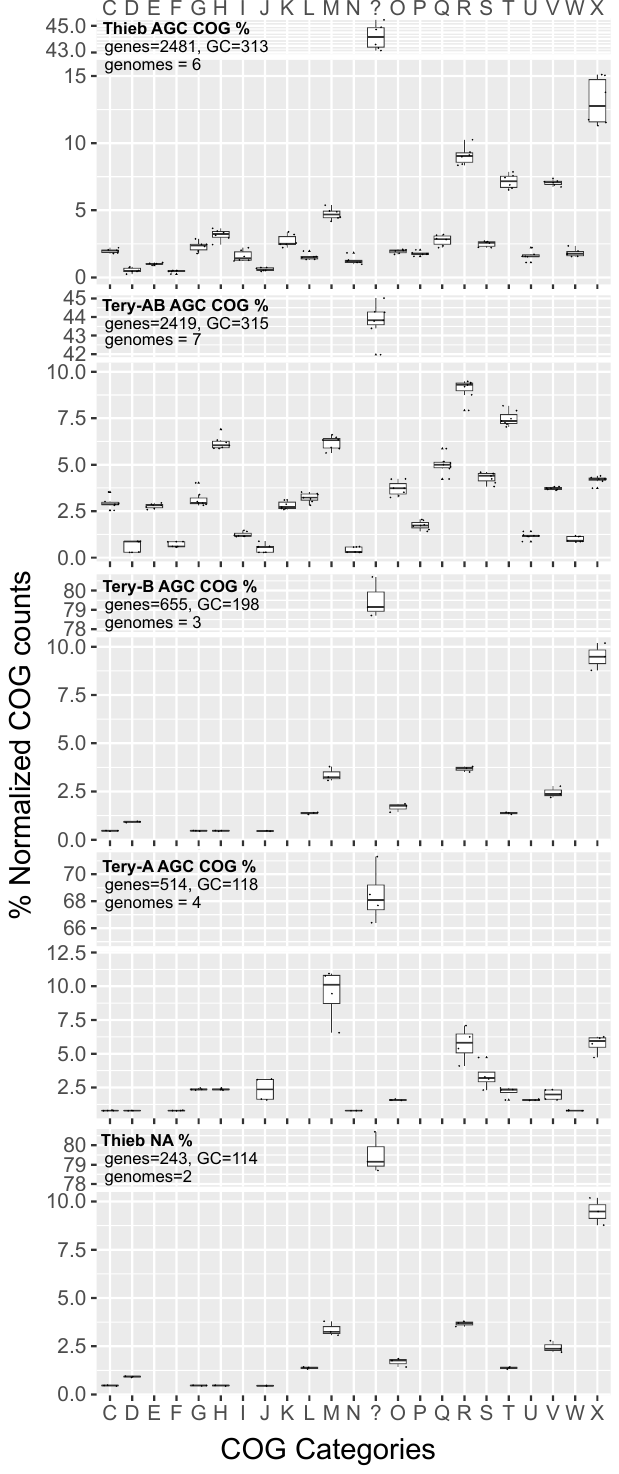
**Supplemental Figure 1.** Percent normalized COG category analysis of AGCs defined in **Fig. 3A.** COG categories are single code abbreviations - J Translation, ribosomal structure and biogenesis, K Transcription, L Replication, recombination and repair, D Cell cycle control, cell division, chromosome partitioning, V Defense mechanisms, T Signal transduction mechanisms, M Cell wall/membrane/envelope biogenesis, N Cell motility, W Extracellular structures, U Intracellular trafficking, secretion, and vesicular transport, O Posttranslational modification, protein turnover, chaperones, X Mobilome, prophage transposon , C Energy production and conversion, G Carbohydrate transport and metabolism, E Amino acid transport and metabolism, F Nucleotide transport and metabolism, H Coenzyme transport and metabolism, I Lipid transport and metabolism, P Inorganic ion transport and metabolism, Q Secondary metabolites biosynthesis, transport and catabolism, R General function, S Function unknown, ? non-categorized.

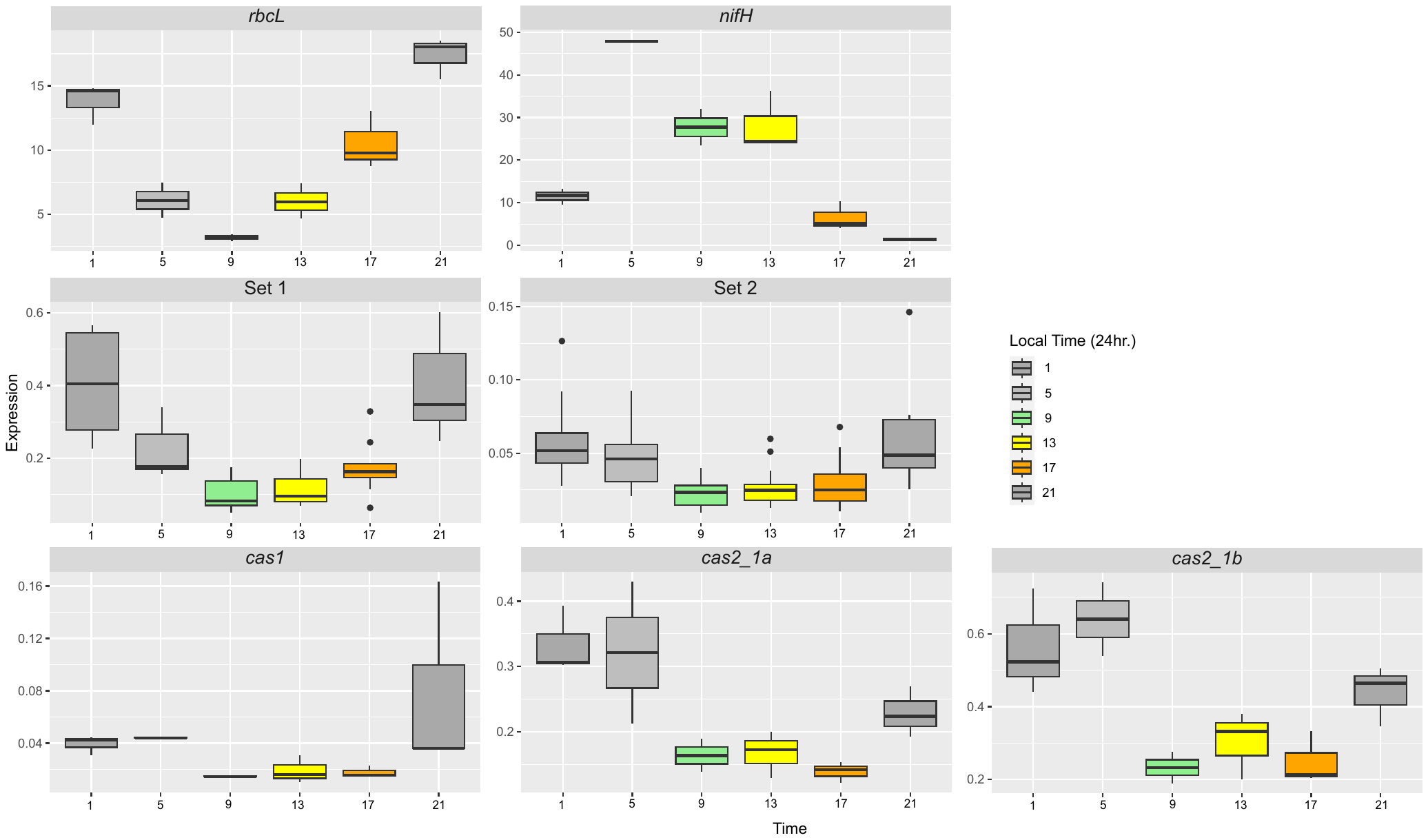

**Supplemental Figure 2.** Normalized read recruiting from North Pacific *Trichodesmium* colony transcripts (88) to H94 *T. thiebautii* *cas* genes shown in Figure 4A compared to key metabolic genes for RuBisCO (*rbcL*) and nitrogenase (*nifH*). Sets 1 and 2 shows the summation of the expression of all *cas* genes in each cluster. Expression for *cas1* and 2 genes are not included in the sets because they are not contiguous with Genes included are *cas10*, *cmr3*, *cmr4*, *cmr6*. All expression data were normalized by a *Trichodesmium* single copy constitutive gene *rotA* (116).
